# Structure-based identification of novel histone deacetylase 4 (HDAC4) inhibitors

**DOI:** 10.1101/2022.05.31.494169

**Authors:** Rupesh Agarwal, Pawat Pattarawat, Michael R. Duff, Hwa-Chain Robert Wang, Jerome Baudry, Jeremy C. Smith

## Abstract

Histone deacetylases (HDACs) are important cancer drug targets. Existing FDA-approved drugs target the catalytic pocket of HDACs, which is conserved across subfamilies (classes) of HDAC. Here, we use molecular modeling approaches to identify and target potential novel pockets specific to Class IIA HDAC-HDAC4 at the interface between HDAC4 and the NCOR protein. These pockets were then targeted using an ensemble docking approach combined with consensus scoring to identify compounds with a different mechanism of binding than the currently known HDAC modulators. Using this approach, 18 compounds predicted in silico to bind to HDAC4’s novel pockets were tested *in vivo* testing on two cancer cell lines. Of these, 5 compounds decreased cell viability to less than 60%. One inhibited the catalytic activity of HDAC4 but not HDAC3, which belongs to a different family of HDACs (Class I). The most potent compound has an IC_50_ comparable to the FDA-approved compound SAHA (Vorinostat). While there are currently no known inhibitors reported to bind highly selectively to HDAC4, the present result suggests potential mechanistic and chemical approaches for the development of selective HDAC4 modulators.

## Introduction

Histone deacetylases (HDACs) catalyze the deacetylation of histone tail lysines, resulting in compaction of DNA and suppression of transcription^1^. HDACs work in opposition to histone acetyltransferase (HAT), which transfers an acetyl group to lysines in the histone tail^2^. In a healthy cell, the transcription levels regulated by HAT and HDAC are well balanced^1^. However, abnormality in the function of HDACs contributes to the initiation and progression of several cancers^1^ and can cause abnormal transcription of critical genes that control vital cell functions, such as proliferation, cell cycle regulation, and apoptosis^3, 4^. HDACs also influence other important genomic functions such as DNA repair, chromatin assembly, and recombination^5^.

HDACs have been assigned to five major classes based on their catalytic mechanisms and sequence homology. Classes I (HDAC 1, 2, 3, 8), IIa (HDAC 4, 5, 7, 9), IIb (HDAC 6, 10) and IV (HDAC 11)^6^ HDACs have a zinc (Zn^+2^) ion in the catalytic pocket as a cofactor in enzymatic activity. In contrast, Class III HDACs, or sirtuins,^7^ require nicotine adenine dinucleotide as a cofactor. HDACs cannot bind to DNA by themselves and exist as components in various multiprotein complexes. Structures of the catalytic domains of all Class I HDACs show only minor differences between them.

Unlike Class I HDACs, Class IIa proteins lack intrinsic deacetylase activity because a key tyrosine residue, which stabilizes the tetrahedral intermediate during deacetylation of the native substrate (acetylated lysine) is absent^8^. The catalytic domain contains two zincs: one structural and one catalytic. A structural zinc is one of the features distinguishing Class IIa from Class I HDACs. However, it is believed that Class IIa HDACs, especially HDAC4, function by forming multiprotein complexes with other enzymatically active Class I HDACs. HDAC4 forms a complex with HDAC3-Nuclear Receptor Corepressor (NCoR) and is believed to operate as a scaffold for the recruitment of multiprotein complexes, thus increasing the range of deacetylation in specific regions of chromatin^9^.

HDAC inhibitors (HDACi) have emerged to be important drugs for cancer therapy, as they can inhibit upregulated HDACs, thus reducing histone deacetylation and leading to normal levels of acetylated histones^10^. To date, four HDACi: Vorinostat (SAHA), Belinostat (PXD101), Panobinostat (LBH589), Romidepsin (FK228) have been approved by the US FDA, and one HDACi, namely Chidamide (CS005) by the Chinese FDA, for the treatments of hematologic malignancies, and more than 20 inhibitors are in clinical trials ^11, 12^. All of these US FDA-approved inhibitors target the HDAC catalytic pocket and share common structural features, including a Zinc-binding group (ZBG), a linker region, and a surface recognition domain^12, 13^. Based on the ZBG chemical structures, HDACis can be divided into five main classes: hydroxamates, cyclic peptides, benzamides, short-chain fatty acids, ketones, and others. However, most hydroxamates are pan-HDACis, while the benzamides have increased Class I selectivity. Three (Vorinostat, Belinostat, Panobinostat) out of the four US FDA-approved drugs are hydroxamate inhibitors that target the evolutionarily-conserved catalytic pocket, thus explaining why these inhibitors are nonspecific binders and potentially toxic^12^.

HDAC4 has been implicated in promoting tumor growth through the suppression of p21 expression in colon cancer, glioblastoma, ovarian cancer, and gastric cancer cells^14^ and therefore is a potential drug target for anti-cancer therapy. However, there are no known inhibitors that target HDAC4 selectively. For the structure-based design of specific inhibitors, there is a need to focus on the structural differences between HDACs ^12, 15^. Here, we aimed to use structure-based drug discovery to identify novel inhibitors that have the potential to selectively bind to HDAC4. Structure-based design is possible since there are two reported crystal structures of the HDAC4 catalytic domain, in open and closed conformations^8^. In the structure of the open conformation a small molecule pan inhibitor is bound in the catalytic pocket, whereas in the closed conformation there is a gain of function mutation of H332Y. The region bound to the structural zinc is closer to the active site in the closed conformation

Most of the enzymatic activity associated with HDAC4 expressed in mammalian cells is due to endogenous HDAC3. HDAC4 can shuttle between the cytoplasm and the nucleus and bind to NCoR, which is also bound to enzymatically active HDAC3^16^. It has been previously reported that HDAC4 binds to repression domain 3 of NCoR, whereas HDAC3 binds to the SANT domain of NCoR^16, 17^. There are no structures of the HDAC4-NCoR complex, but based on mutational studies ^18^ 21 “hot spot” residues (C667, C669, C751, D759, T760, S767, A774, P799, P800, G801, H803, A804, F812, C813, H842, H843, G844, N845, G846, G868, and F871) present on the surface of HDAC4 have been identified as preventing binding to the NCoR-HDAC3 complex in vivo (**Figure 1**). These interfacial residues are conserved across all Class IIa HDACs and include the residues in the opening of the catalytic pocket. In contrast, some of these residues (C667, C669, C751, D759, T760, and F871) are only found in Class IIa HDACs and not in Class I, which potentially forms the basis for interaction specificity with NCoR.

**Figure 1:**
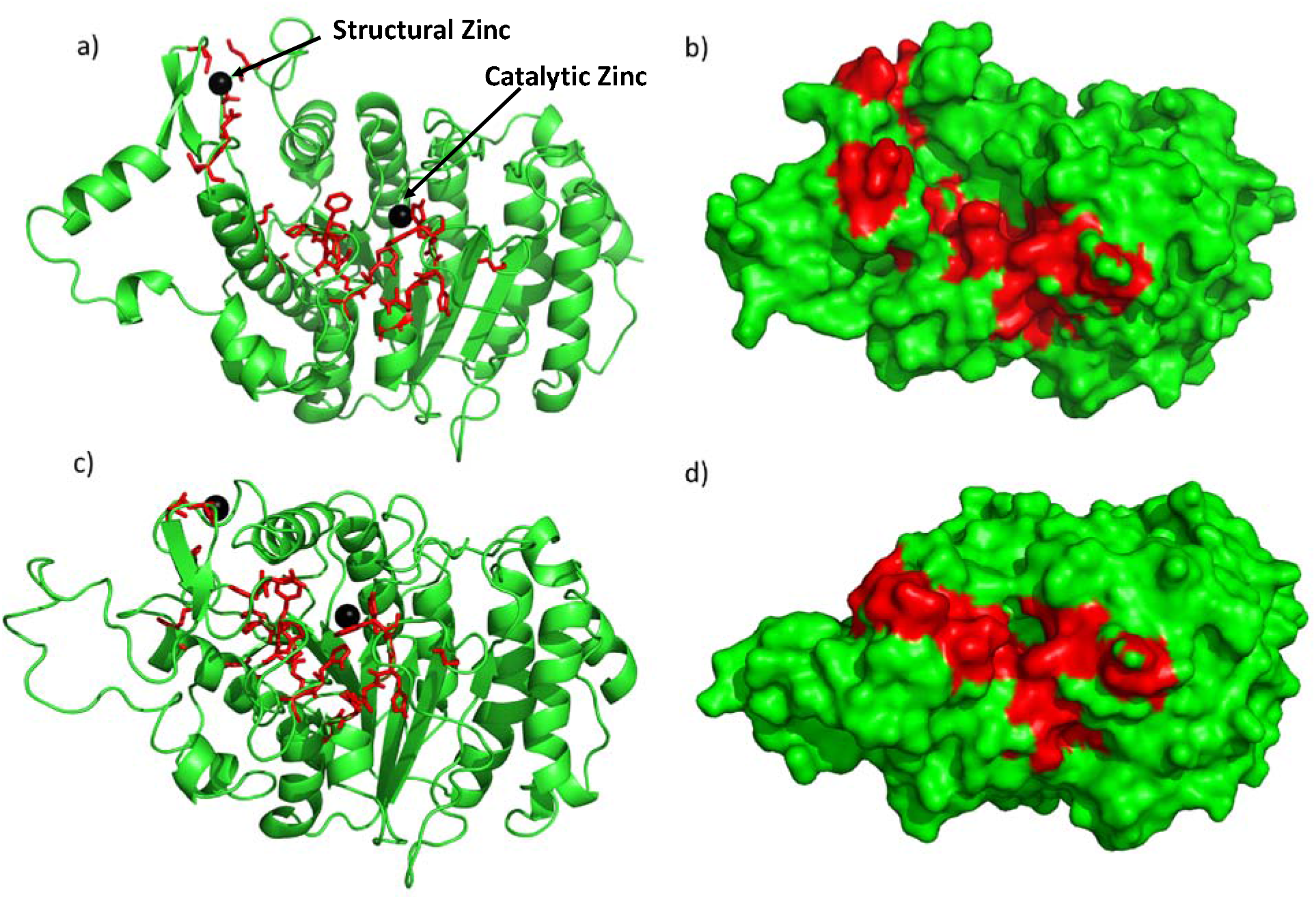
Crystal structure of the open conformation^8^ (a & b) and the closed conformation^8^ (c & d) of HDAC4. The red sticks and red region represent residues classified as interfacial “hot spot” residues identified previously^18^; black spheres represent the two zincs (catalytic & structural). a & c are ribbon models of the crystal structures, and b & d are surface models.

In an attempt to engineer some level of selectivity our strategy focused on the HDAC4 interface with NCoR rather than targeting the conserved catalytic pocket. Protein-protein interfaces (PPIs) have emerged as major drug targets since a large number of proteins are critical in biological pathways related to various diseases function after complex formation^19^. However, there are major challenges in targeting PPI interfaces since they often lack cavities for small molecules to bind or are intrinsically disordered^20, 21^. Nevertheless, small molecules interacting at interface “hot-spots” can compete with the binding of the cognate partner without necessarily covering the whole PPI surface^22-24^.

We used classical molecular dynamics (MD) and enhanced sampling accelerated molecular dynamics (aMD) simulation^25^ approaches to generate an ensemble of structures and to identify potential pockets in the experimentally characterized interface of HDAC4 and NCoR. We thus identified novel pockets in the PPI which are not present in the crystal structures. We hypothesized that small molecules binding in these pockets could lead to disruption of complex formation. We performed high throughput *in silico* screening of small molecule libraries on the ensemble of structures generated from the simulation. Using this ensemble docking approach^26^, we identified 18 compounds that were tested in cell viability assays, five of which decreased cell viability to less than 60%. One of these compounds inhibited the catalytic activity of HDAC4 but not HDAC3 while, serendipitously, another inhibited the catalytic activity of HDAC3 but not HDAC4.

## Methods

### System preparation

Crystal structures of HDAC4 in the open (PDB 2VQM)^8^ and in the closed state (PDB 2VQW)^8^ were used for this study. The protein structures were modeled using Amber ff14SB parameters^27, 28^ and water was modeled as TIP3P^29^. Using xleap, cysteines 667, 669, and 751 were deprotonated (-ve charge) and converted to CYM for the closed conformation, and cysteines 667 and 751 were converted to CYM for the open conformation as observed in the respective crystal structure. The open structure was crystallized in the presence of an inhibitor, which was deleted for the simulation. The closed structure was crystallized with a mutation H332Y, and so the tyrosine 332 was mutated to wild type histidine for the simulation. The protein was surrounded by an octahedral box of TIP3P^29^ of 10 Å around the protein in each direction.

### Classical molecular dynamics simulation

MD simulations were performed on both crystal structures. First, the HDAC4 structure was held fixed with force constant of 500 kcal mol^-1^ Å^-2^ while the system was minimized with 500 steps of steepest descent followed by 500 steps with the conjugate gradient method. In the second minimization step, the restraints on HDAC4 were removed, and 1000 steps of steepest descent minimization were performed, followed by 1500 steps of a conjugate gradient. The system was heated to 300 K while holding the protein fixed with a force constant of 10 kcal mol^-1^ Å^-2^ for 1000 steps. Then, the restraints were released, and 1000 MD steps were performed. The SHAKE algorithm^30^ was used to constrain all bonds involving hydrogen in the simulations. A 50 ns MD was performed at 300 K using the NPT ensemble and a 2 fs time step. The temperature was fixed with the Langevin dynamics thermostat^31^ and the pressure was fixed with a Monte Carlo barostat^32^. This procedure yielded a total of 250,000 snapshots for subsequent analyses.

### Accelerated molecular dynamics simulation

Accelerated molecular dynamics (aMD)^25^ samples conformational space more efficiently than classical MD, resulting in receptor conformers which would not have been otherwise sampled. Unlike other enhanced sampling methods, aMD does not require reaction coordinates or collective variables and it is well suited for creating an ensemble of structures to characterize the conformational flexibility of a receptor. The aMD simulation was performed by using a dual energy boost (i.e. dihedral energy and potential energy) with an acceleration parameter (α) of 0.2. The average total potential energy and average dihedral energy parameters for aMD were calculated from the classical MD simulation. We performed 100 ns of aMD simulation. This procedure yielded a total of 500,000 snapshots for subsequent analyses. Snapshots from classical MD and aMD were clustered by root mean square deviation (RMSD) of hot spot residues with the hierarchical agglomerate clustering algorithm present in the Cpptraj module^33^.

### *In silico* screening

We generated an ensemble of 4 representative snapshots from each of the classical MD and aMD clusters for both the open and closed conformations. The NCI Diversity V set (∼1600 compounds)^34^ was docked to a total of 9 receptor conformations (4 snapshots from MD, 4 from aMD and 1 crystal structure) with a cubic box size of ∼30 Å. VinaMPI^35^, a parallel version of AutodockVina^36^, was used to perform the *in silico* screening. The docked poses were then ranked by the AutodockVina scoring function^37^. These docked poses were then refined using the ‘induced fit’ tool in MOE^38^ in their respective pockets. The poses of the top 100 compounds were manually checked (to maximize the pocket coverage, maximize the structural diversity of the compound, checked for the type of interaction (electrostatic or hydrophobic), and molecule availability at NCI) and 18 compounds were suggested for experimental testing.

### Cell viability assay

Cell viability assays were determined using Methyl Thiazolyl Tetrazolium (MTT) assay (Travigen, Gaithersburg, MD, USA). Five × 10^3^ SW780 (human transitional urinary bladder cancer cell line) or 1 × 10^4^ (human triple negative breast cancer cell line MDA-MB231, American Type Culture Collection [ATCC], Rockville, MD) in 100 µL of culture media were seeded to each well of 96-well culture plates. MDA-MB231 and SW780 cell line were cultured in Dulbecco’s modified Eagle’s medium (DMEM) with 10% or 5% heat-inactivated fetal bovine serum, respectively, 100□U/mL penicillin, and 100□µg/mL streptomycin. Cultures were maintained in 5% CO_2_ at 37□°C and sub-cultured every 2–3 days. At 24 hours after seeding, cells were treated with various concentrations of potential HDAC inhibiting compounds for 48 hours (100 µM for initial screening, 0.01, 0.01, 1, 10, 100, 250, and 500 µM for dose-response experiment). Subsequently,10 µL of MTT reagent were added to each well and incubated for 4 hours, followed by addition of 100 µL of detergent reagent. The plate was incubated overnight. The reduced MTT reagent in cultures was measured at 570 nm by using an ELISA reader (Bio-Tek, Winooski, VT, USA). The IC50 for each compound was calculated from fitted non-linear regression dose-response curves.

### HDAC activity assay

HDAC3 and HDAC4 activity assays were performed using fluorogenic assay kits obtained from BPS Bioscience (San Diego, CA) by following the protocol that is supplied by the manufacturer. Briefly, purified enzyme (HDAC3 or HDAC4) in buffer containing 0.1% BSA was incubated with substrate at 37 °C for 30 minutes in a 96-well plate. Fluorescence developer was added and the plate incubated at room temperature for a further 15 minutes. The activity of the enzymes in the absence and presence of compounds was measured on a BioTek Cytation 5 plate reader, exciting the samples at 350 nm and measuring emission at 450 nm. Compounds were prepared as 10 mM stocks in DMSO, diluted to 1 mM in 20 mM NaHEPES, 150 mM NaCl, pH 7.4 buffer, and were screened at 100 μM in the assays at least in duplicate. A positive control with 1% DMSO added in lieu of inhibitor and a negative control without enzyme were also prepared in each assay plate. The negative control was subtracted from all fluorescence values, while the positive control was used as the reference for 100% activity of the enzyme. IC50 values were determined for the compounds that decreased HDAC activity by at least 25% relative to the no-inhibitor positive control.

## Results

### Novel pockets from MD simulations and ensemble docking

The PPI residues on HDAC4 encompass the catalytic pocket and structural zinc and the region between them. In the crystal structures, this region is rather smooth and has no obviously druggable cavity for a small molecule to bind, which is a common drawback in targeting PPIs^20, 21^. However, in the ensemble of receptor structures generated from classical MD and aMD simulations for both open and closed conformations, we identified novel cavities/pockets in the PPI which were not present in the crystal structures (**Figure 2**). These cavities are present in the “hot-spot” region and provide potential pockets for small molecules to bind. These pockets were generated by clustering of the trajectory, and hence are present for significant proportions of the simulation time. Therefore, they are low energy states and not rare conformations. We observed some shallow cavities (**Figure 2**), and a deep pocket near the structural zinc, which contains PPI hot spot residues as shown in **Figure 2**. These pockets stretch throughout the PPI region. These results illustrate the usefulness of generating an ensemble of structures rather than using only the crystal structure.

**Figure 2:**
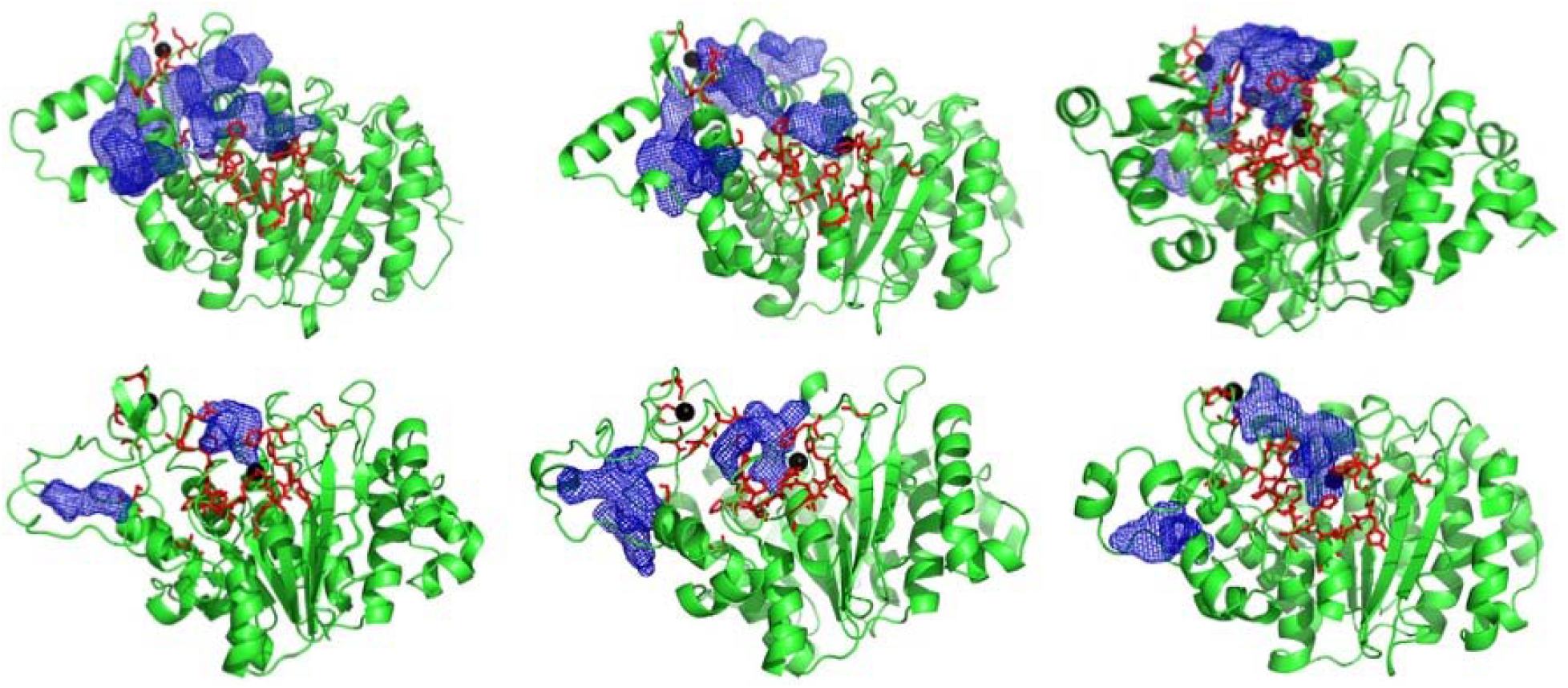
HDAC4 ensemble structures showing novel pockets/cavities (blue mesh) generated from simulations of crystal structures. Red sticks represent PPI “hot spot” residues identified experimentally.

The docking search box was set to be large enough to include both the catalytic pocket and the novel pockets identified, as both are in the PPI region. Since the catalytic pocket itself is large and hydrophobic, we found that many of the compounds ended up in the catalytic pocket. Therefore, we filtered out those compounds which were within 10 Å of the catalytic zinc. The aim of this approach was to identify compounds that bind with a high score in the PPI. We found small molecules bound in the novel pockets discovered (**Figure 3**). These docked poses were further refined in their respective pockets using the ‘induced fit’ tool in MOE^38^ to further improve the hit rate^39^. After manually examining the docked poses, we picked 18 high-scoring compounds chosen to maximize the pocket coverage (**Figure S2**), for chemical diversity (**Figure S1**), and for the availability of compounds at NCI. These compounds were then suggested for experimental testing.

**Figure 3:**
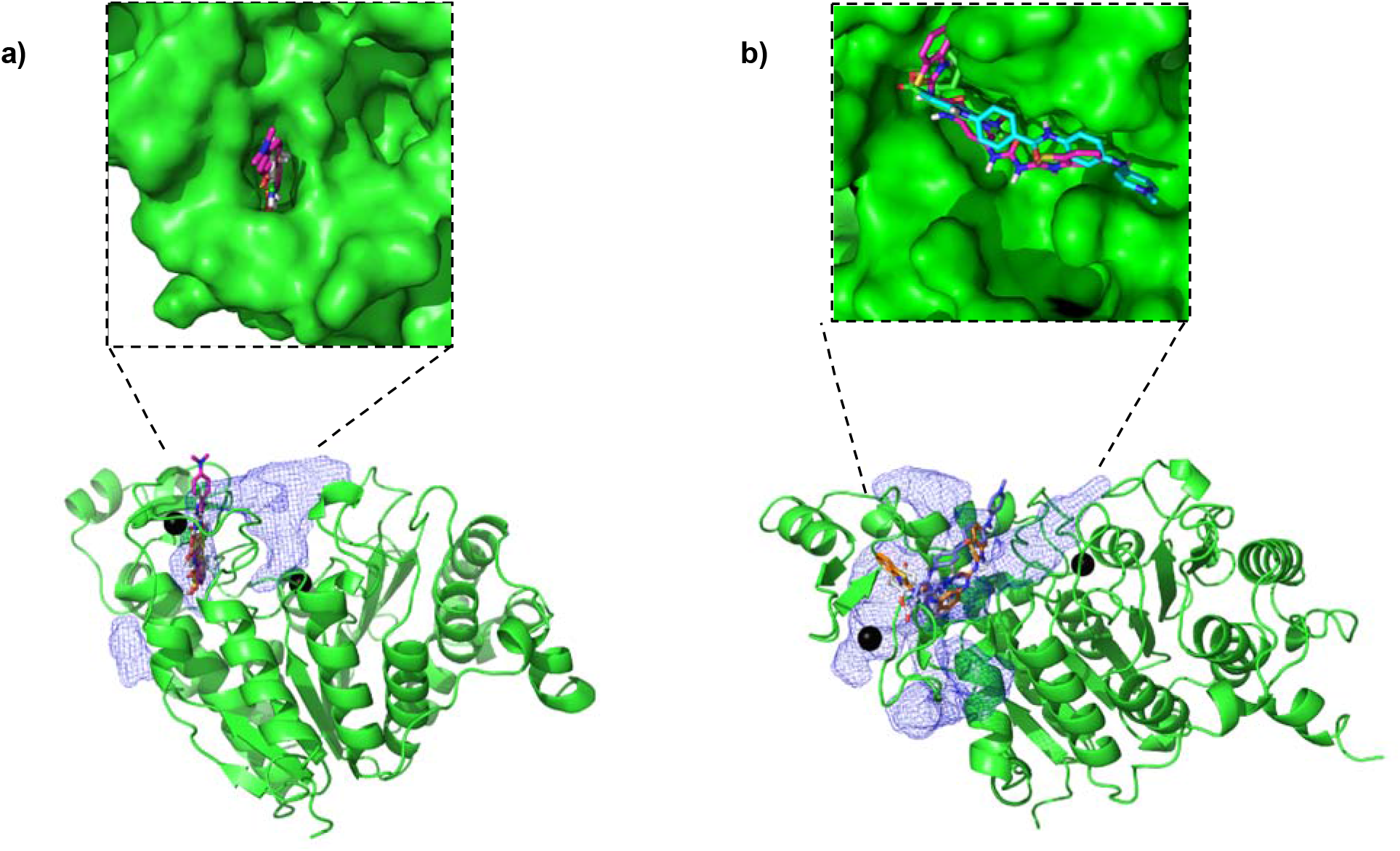
Some of the top ranked compounds bound in the novel pockets identified in a) closed conformation and b) open conformation ensemble structure

### Cell susceptibility to identified small molecules

To test the in silico derived compounds in vivo, we selected 18 NCI compounds (NCI IDs: 117922, 134199, 319435, 34488, 36425, 373535, 4135, 51936, 55172, 60034, 79887, 88402, 42231, 11926, 195327, 299968, 44584, 67436) for testing on human breast carcinoma MDA-MB231 and transitional urothelial carcinoma SW780 cell lines. Both these cell lines have been shown to significantly upregulate HDACs^40, 41^. These 18 compounds were tested in a cell viability assay at 100 µM, with DMSO as a negative control and SAHA (Vorinostat, a US FDA-approved drug)^42^ as a positive control.

5 compounds: 67436, 11926, 299968, 44584 and 195327, were able to reduce cell viability of MDA-MB231 to lower than 60% (**Figure 4a**) and were considered as active. Moreover, 67436, 11926, and 195327 were also able to reduce SW780 to lower than 50%, indicating that these three compounds are effective against both cancer cell lines. Next, compounds (195327, 44548, 299968, 67436, 11926) that reduced cell viability to less than 60% were further examined by dose-response exposure of each compound at 0.01, 0.01, 1, 10, 100, 250, and 500 µM, and the IC_50_ of each compound calculated. A concentration-dependent decrease of cell viability was observed for all the selected compounds (**Figure 4b**). The most potent compound is 195327, which has an IC_50_ comparable to SAHA at 37±17 µM vs. 35±10 µM for MDA-MB231 and 89±48 vs. 90±45 for the SW780 cell line. Data also showed that 299968 and 44584 were effective against SW780 but at higher doses than SAHA (IC50 = 300±17 and 213±13, respectively). Compound 11926 reduced cell viability of both MDA-MB231 and SW780, but the efficacy against SW780 was lower than against MDA-MB231(Cell viability = 45% at 500 µM in SW780, compared to 9% at 500 µM in MDA-MB231). These results indicated that the compounds are effective against both cancer cell lines, but with different levels of response.

**Figure 4.**
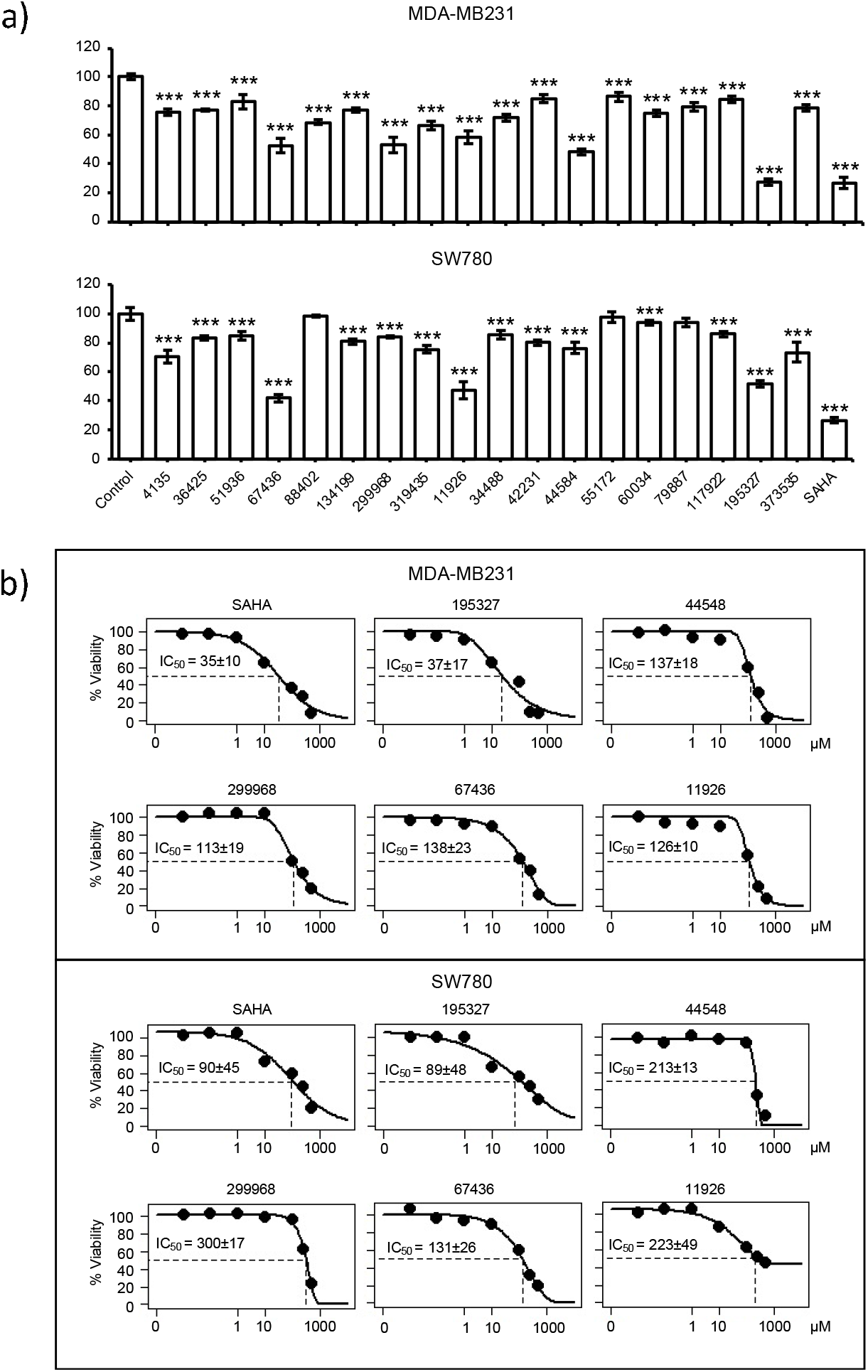
Effect of compounds on cell viability of cancer cell lines. a) Relative cell viability of MDA-MB231 and SW780 cell lines 48 hours after treatment with candidate compounds determined by MTT assay. Relative cell viability was normalized by the value determined in untreated counterpart cells set as 100%. Columns, mean of triplicates; bars, SD. Statistical significance was determined using student’s t-test (a, e, & f), indicated by ***p < 0.001. b) Dose-response exposure of compounds that reduced cell viability to less than 60%. IC_50_ of each compound was calculated.

### HDAC Activity

With the results from the cell viability assays, we decided to dig deeper and performed enzyme assays. Since the cell lines used also contain upregulated Class I HDACs, the 18 compounds were screened for their ability to inhibit HDAC4 and HDAC3 (from Class I HDAC subfamily) (**Figure 5**) *in vitro*. At 100 μM, two compounds (67436 and 134199) decreased both HDAC3 and HDAC4 activities to 40% or less, and a number of other compounds were specific to one of the enzymes. HDAC4 was significantly inhibited by 88402 (<40% activity retained), but the compound only mildly perturbed HDAC3 activity (75% activity); however, that compound did not show activity in the in-vivo assay (**Figure 4a**). Compounds 34488, 44584, and in vivo active compound 195327 decreased HDAC3 activity by 50%, 60%, and 80%, respectively, whereas HDAC4 activity remained unchanged in the presence of those compounds (**Figure 5**).

**Figure 5.**
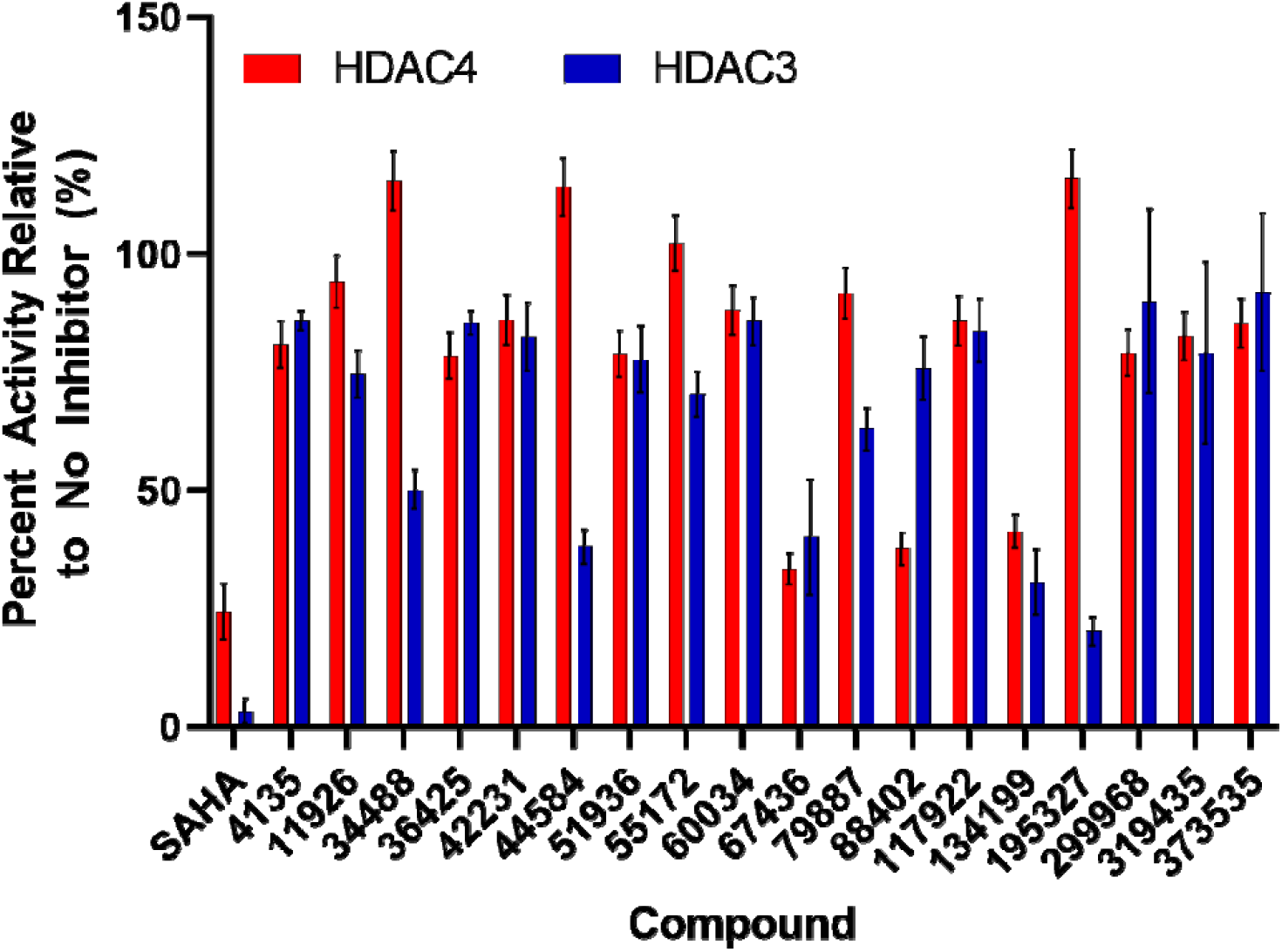
Effect of compounds on the activities of HDACs. The relative percent activity of HDAC3 (blue bars) and HDAC4 (red bars) in the presence of 100 μM inhibitor compound compared to the absence of inhibitor. Error bars are the standard deviation of at least two assays.

The compounds that decreased the activity of HDAC4 by at least 25% were further screened for IC50 values (**Table 1, Figure S3, S4, S5**). Compounds 67436 and 134199 had IC50 values in the low micromolar range for both HDACs, which is again comparable to the known inhibitor of HDAC4, SAHA. However, both compounds were significantly weaker inhibitors of HDAC3 than SAHA and thus more selective, albeit to a limited extend. Compounds 88402, 319435, and in vivo active compound 299968 had IC50 values at least 2-fold lower for HDAC4 than HDAC3, showing that these compounds are potentially also HDAC4-specific inhibitors. Both 299968 and 319435 decreased the activity of HDAC3 and HDAC4 to similar levels in the initial compound screen (**Figure 5)**. However, these compounds did not inhibit HDAC3 up to a concentration of 500 μM in the IC50 experiments, indicating they most likely do not inhibit HDAC3 **(Figure S3, S4, S5**). The higher errors in the screening assay for HDAC3 (**Figure 5**) with compounds 299968 and 319435 may have masked differences in inhibition of the two enzymes by these compounds. The other compounds that weakly inhibited HDAC4 (4135, 36425, and 51936) had IC50 values not much lower than the highest concentration used in the assay (500 μM) and may not be significant inhibitors of either enzyme. Also, while compound 195327 yielded promise as an inhibitor of HDAC3 in the initial compound screen, no inhibition was noted in the IC50 experiments. Also, 195327 did not inhibit HDAC4 in either assay suggesting the inhibition of HDAC3 by the compound noted in the initial screen may have been artefactual. Interestingly, compound 134199 was one of the better inhibitors of both HDACs in the enzyme assay but had little potency in the in vivo screen. From these results, we have identified 8 compounds that bind to HDAC4 or HDAC3 and were able to inhibit the deacetylase activity in the assay. Out of these 8, 3 compounds (88402, 299968, and 319435) bind with some level of weak selectivity to HDAC4 according to IC50 data, and one compound (67436) binds slightly more to HDAC3.

**Table 1.** IC50 values for compounds that inhibited HDAC3 and HDAC4

| **Compound** | **IC<sub>50</sub> (μM) for HDAC3** | **IC<sub>50</sub> (μM) for HDAC4** |
| --- | --- | --- |
| SAHA | 0.024 ± 0.006 | 25±13 |
| 4135 | >500 | 410 ± 240 |
| 11926 | >500 | ND |
| 34488 | >500 | ND |
| 36425 | >500 | 480 ± 260 |
| 44584 | >500 | ND |
| 51936 | >500 | 550 ± 350 |
| 55172 | >500 | ND |
| 67436 | 9.2 ± 5.2 | 47±28 |
| 79887 | >500 | ND |
| 88402 | 210 ± 60 | 100±40 |
| 134199 | 25 ± 7 | 33±12 |
| 195327 | >500 | ND |
| 299968 | >500 | 250 ± 170 |
| 319435 | >500 | 150 ± 50 |

## Discussion

HDACs are cancer drug targets, but most of the FDA-approved HDAC drugs are pan HDAC inhibitors and selectivity towards a specific sub-family is difficult to achieve. Hence, in this study, we moved away from the common strategy of targeting the conserved catalytic pocket and focused on a specific protein-protein interface. HDAC4 complexation with NCoR and HDAC3 is critical for scaffolding function and disrupting the complex can lead to functional inhibition. However, the PPI interfaces present in the crystal structures, like many PPIs, are smooth and do not have obvious cavities/pockets that can be targeted by small molecules. Therefore, we performed classical and enhanced sampling MD simulations to sample different conformations of the PPI. Indeed, we were able to identify pockets in the PPI region that have not been characterized before and are not present in either of the crystal structures. Using ensemble docking, we targeted these novel pockets present in the PPI region and performed an ‘induced fit’ refinement. The aim of this study was to find a novel hit molecule with a novel scaffold that could be used in the future as a precursor to selective inhibitors.

By combining computational and experimental methods, we were able to identify novel compounds that reduce cell viability in human breast carcinoma MDA-MB231 and transitional urothelial carcinoma SW780 cell lines as well as in *in vitro* assays. Out of 18 compounds that were tested, 5 NCI compounds showed a reduction in cell viability to below 50% and have IC50s comparable to the FDA-approved drug SAHA (Vorinostat) but very different scaffolds than both SAHA and the other 4 known drugs when compared using MACCS fingerprinting (**Figures S6, S7, Table S1)**. However, the toxicity profile of these molecules was not tested. In vitro activity assays indicate that one of these (67436) inhibits the activity of both HDAC4 and HDAC3 whereas others had selective inhibition for either HDAC3 or HDAC4.

Dose-dependent activity assays suggested that three compounds, 88402, 299968, and 319435 bind selectively to HDAC4, with IC50s of 100±40 μM, 250 ± 170 μM, and 150 ± 50 μM, respectively, while 67436 was selective for HDAC3 with an IC50 of 9.2 ± 5.2 μM. 195327, 44548, and 11926 showed high activity in cell viability assay but were not as active in the in vitro assay. We hypothesize that these molecules might bind in the distal PPI region of HDAC4 away from the catalytic pocket, which has been previously reported to be critical for HDAC4 function^43, 44^, but this could not be tested using the activity assay.

The results highlight the usefulness of ensemble docking in early-stage drug discovery and identifying molecules with novel chemical matter. Starting with the goal of finding novel HDAC4 modulators, we serendipitously also found one HDAC3 selective modulator. These compounds are offered to the community as potentially good starting points for hit expansion, lead optimization steps, or a degrader (PROTACS) strategy^45^. Additional structural characterization using techniques such as X-ray crystallography and optimization of ADMET (absorption, distribution, metabolism, excretion, and toxicity) properties would be useful in guiding lead optimization and associated efficacy improvement. Moreover, these molecules need to be further tested with other class IIa and I HDACs to understand their selectivity profile and mechanism of action.

## Supporting information

All Supplementary figures and tables

