## Supplementary material for "Structure-based identification of novel histone deacetylase 4 (HDAC4) inhibitors": All Supplementary figures and tables

*Co-corresponding authors

**Supplementary information**

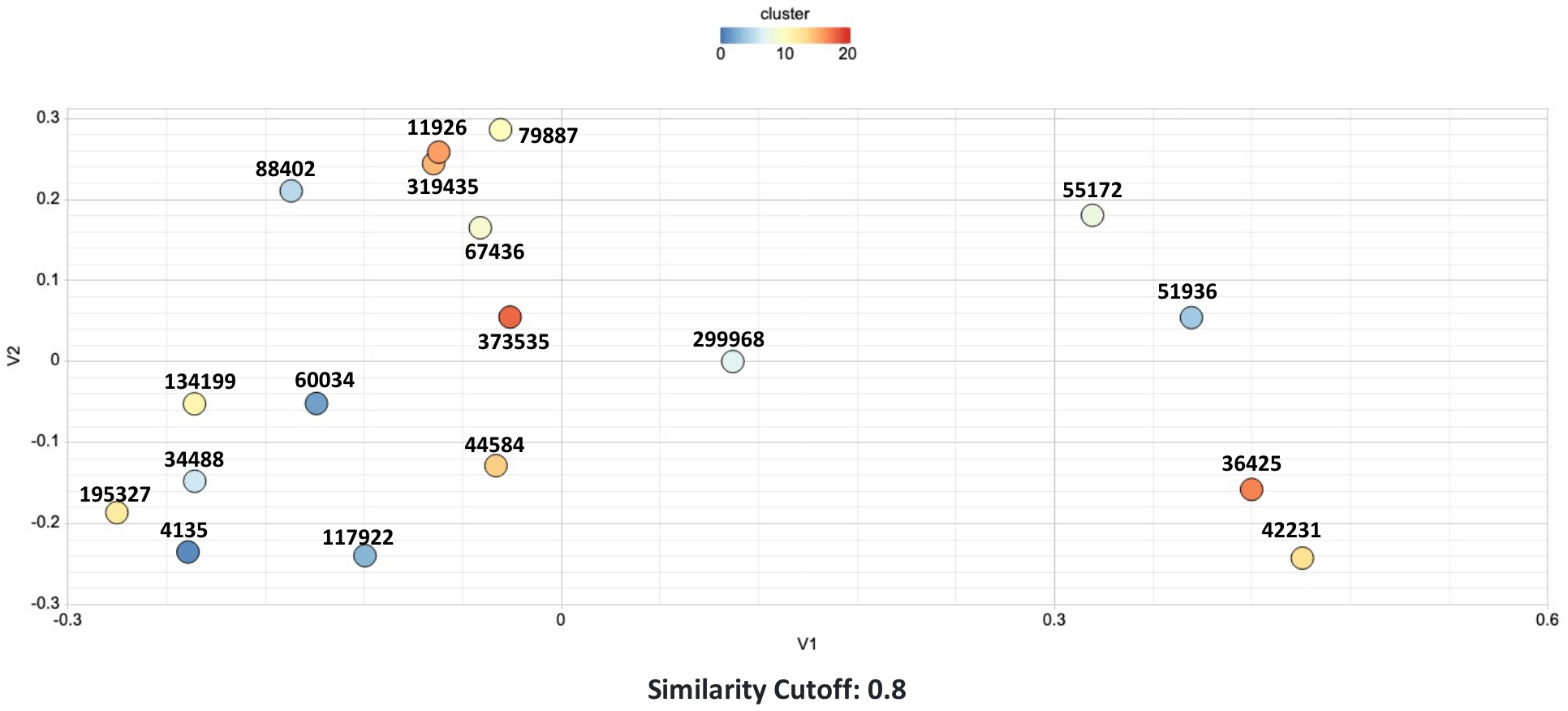

**Figure S1: Clustering of compounds by structural and physicochemical similarities by Multidimensional Scaling (MDS) using ChemmineTools^44^. Similarity cutoff of 0.8 is used for coloring only.**

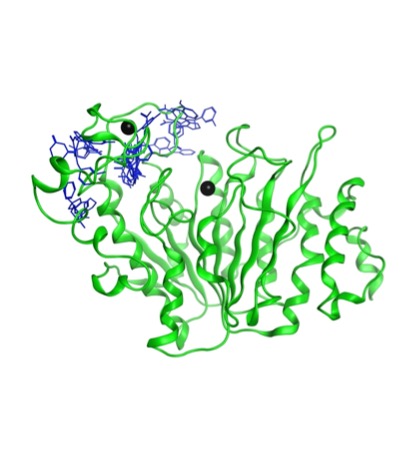

**Figure S2: Some of the top hit molecules (in Blue) and there poses on HDAC4 (in green ribbon) with Zinc atoms (as black sphere) to shown binding site coverage. The figure was made after superimposition of the hits with their respective ensemble structure. All the ensemble structures and hits are not shown for clarity.**

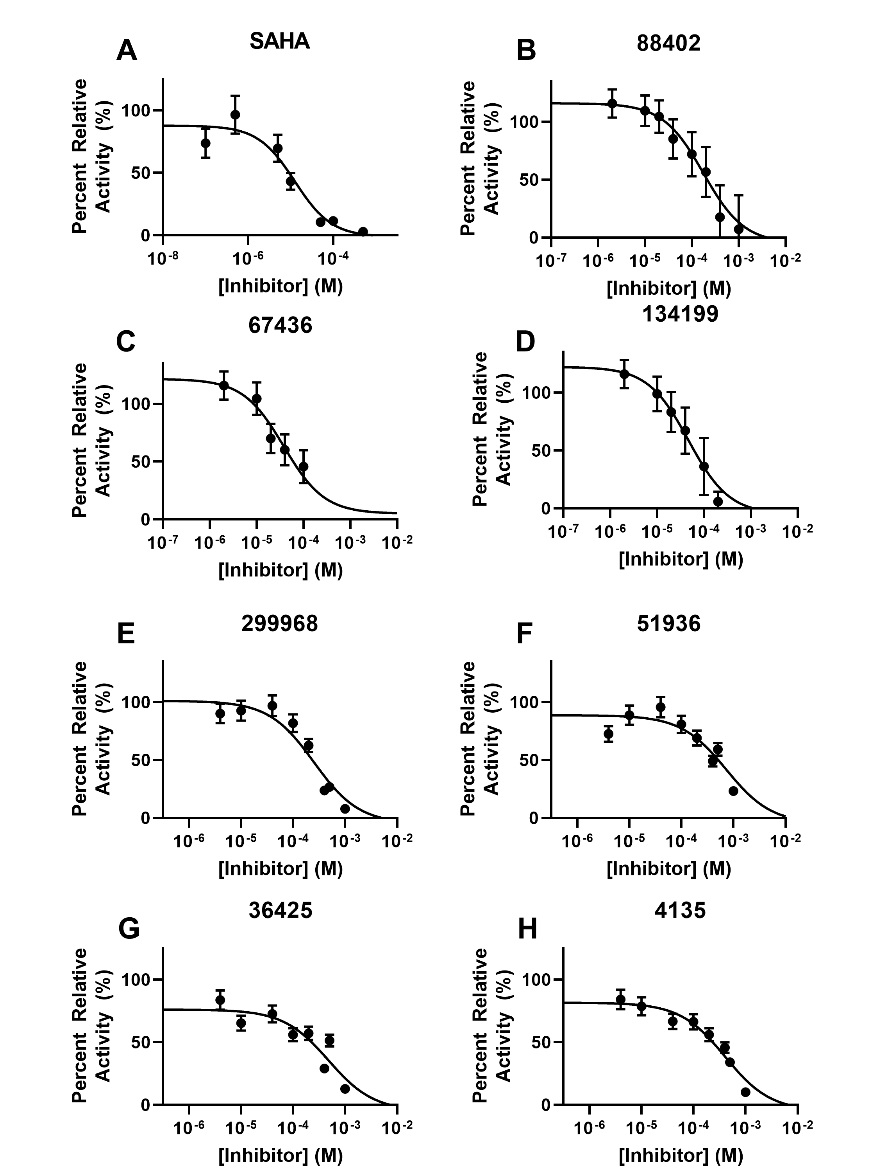

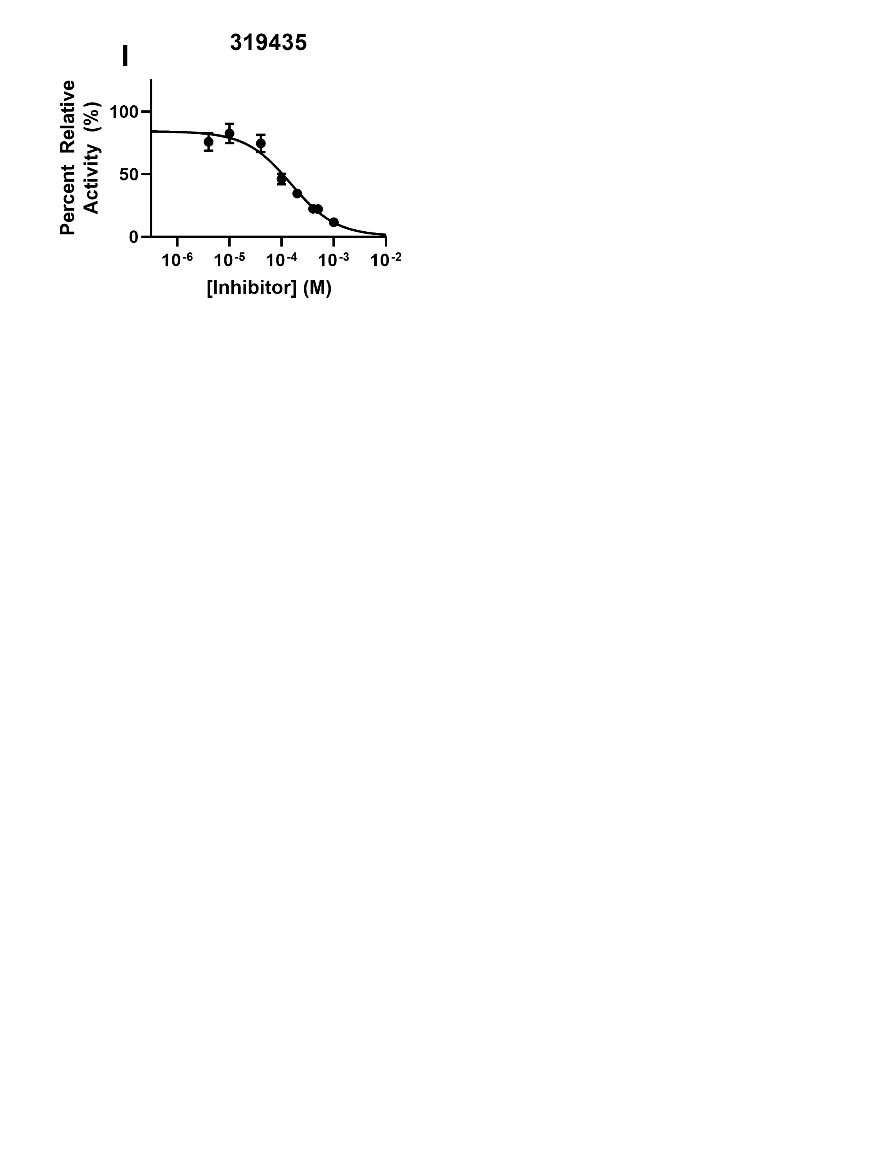

**Figure S3. IC50 plots for compounds that inhibit HDAC4.**

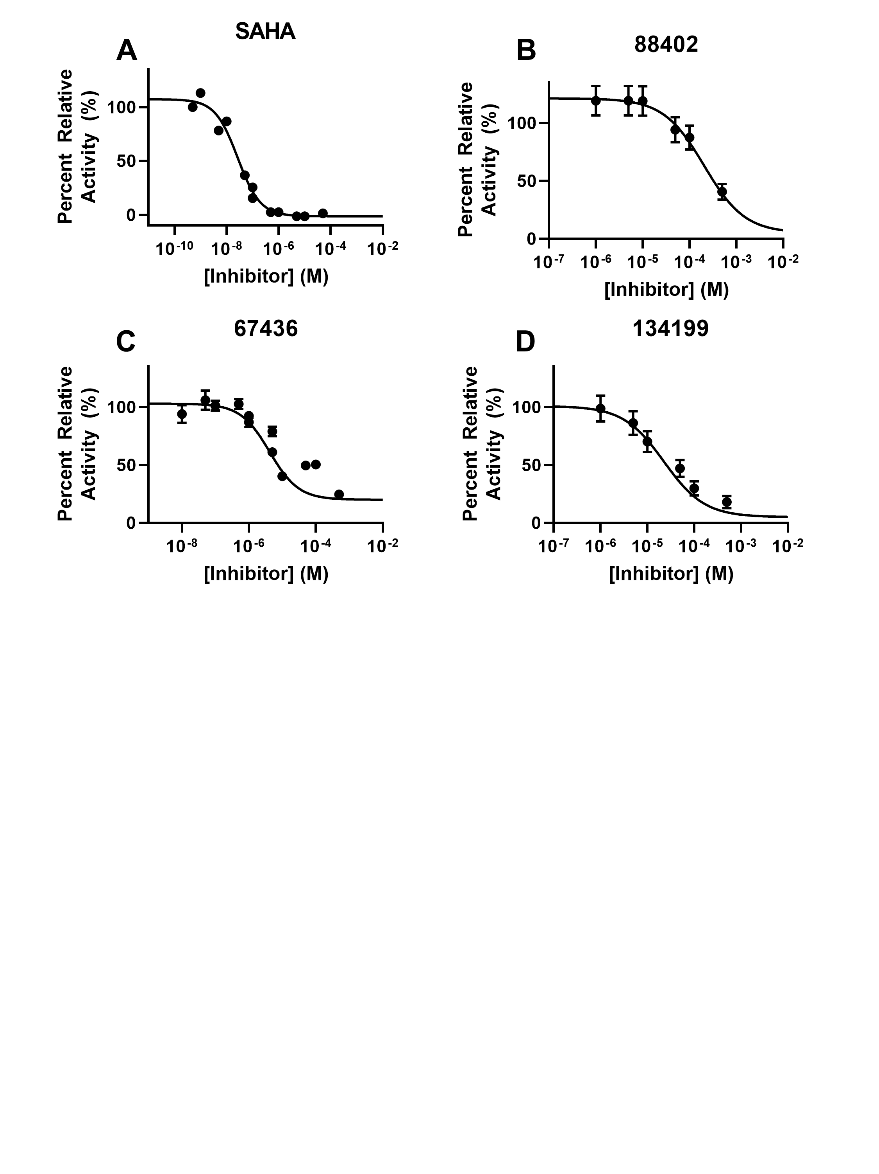

**Figure S4. IC50 plots for compounds that inhibit HDAC3.**

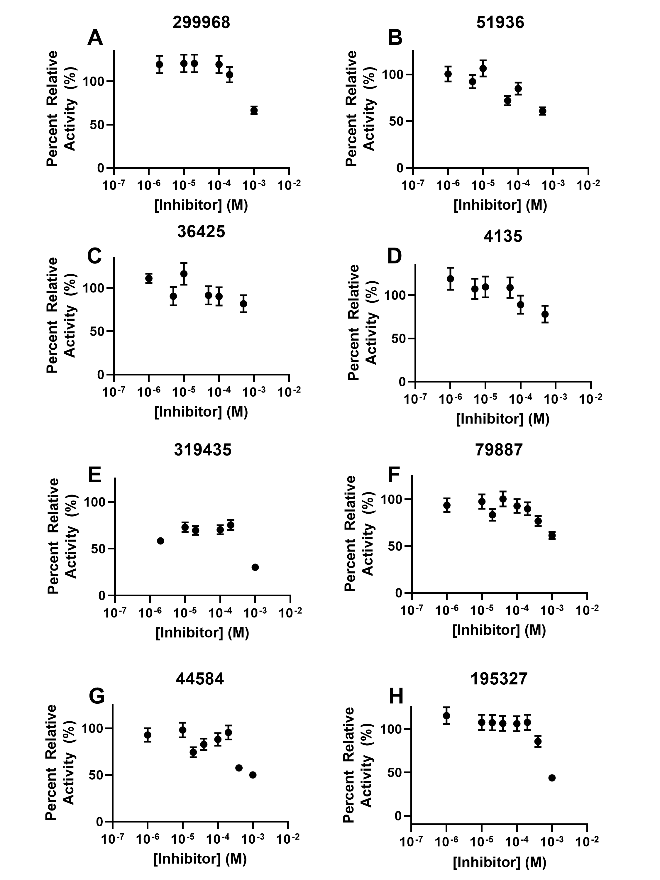

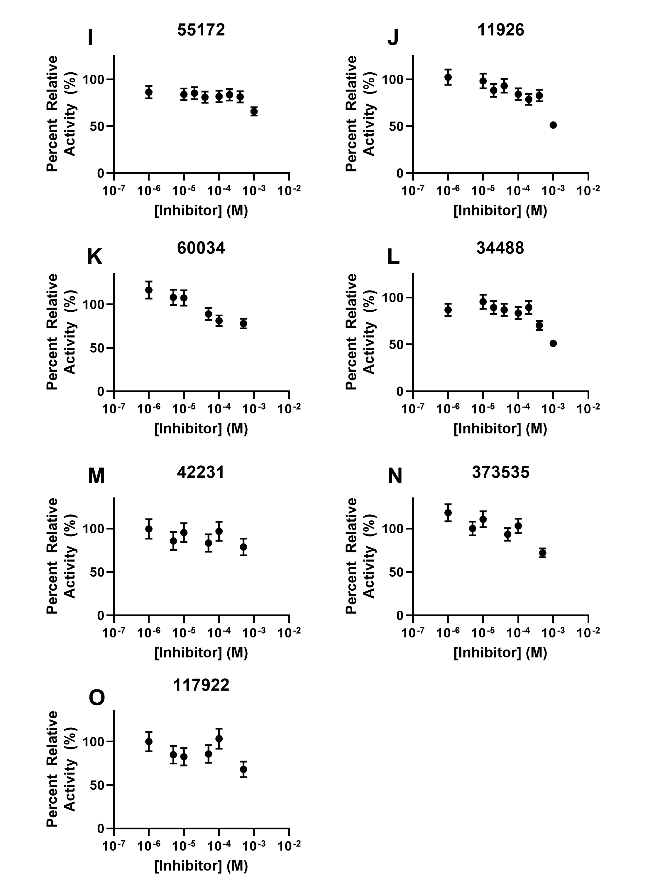

**Figure S5. IC50 plots for compounds that yielded no inhibition of HDAC3 below 500 μM.**

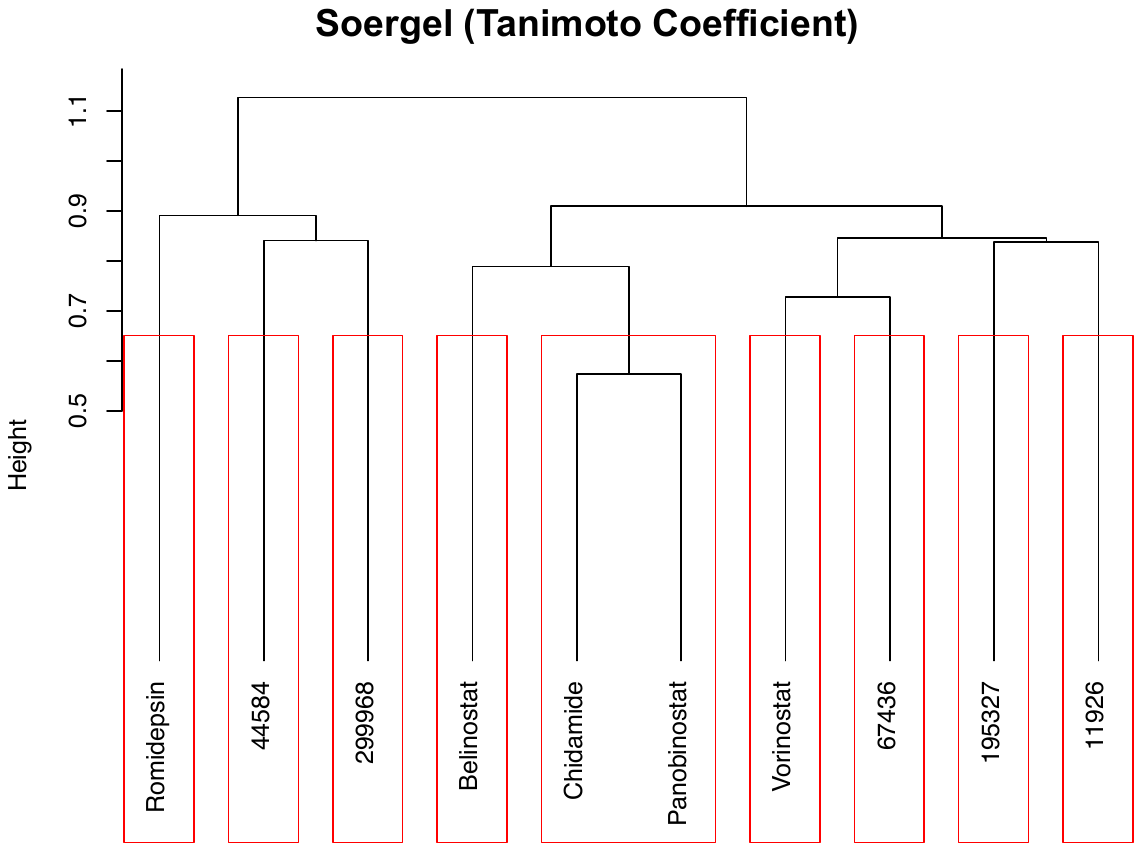

**Figure S6: Hierarchical Clustering using MACCS fingerprinting. Distance Method Soergel (Tanimoto Coefficient); Clustering Method: Ward Linkage; Clustering Threshold: 0.7**

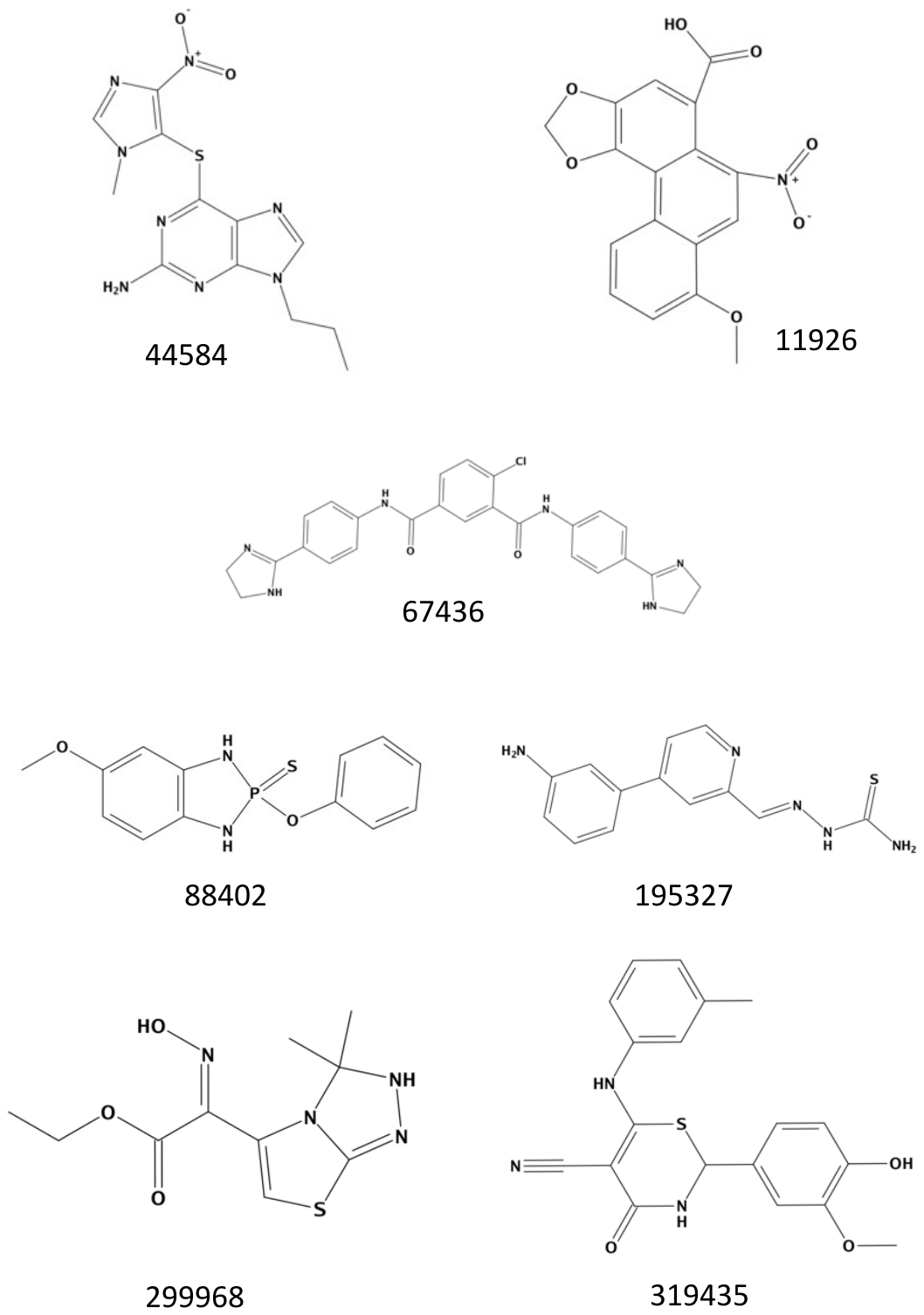

**Figure S7: Two-dimensional structure of top hits.**

| **ID** | **Canonical SMILES** |
| --- | --- |
| 44584 | CCCN1C=NC2=C1N=C(N=C2SC3=C(N=CN3C)[N+](=O)[O-])N |
| 11926 | COC1=CC=CC2=C3C(=C(C=C21)[N+](=O)[O-])C(=CC4=C3OCO4)C(=O)O |
| 67436 | C1CN=C(N1)C2=CC=C(C=C2)NC(=O)C3=CC(=C(C=C3)Cl)C(=O)NC4=CC=C(C=C4)C5=NCCN5 |
| 88402 | COC1=CC2=C(C=C1)NP(=S)(N2)OC3=CC=CC=C3 |
| 195327 | C1=CC(=CC(=C1)N)C2=CC(=NC=C2)C=NNC(=S)N |
| 299968 | CCOC(=O)C(=NO)C1=CSC2=NNC(N12)(C)C |
| 319435 | CC1=CC(=CC=C1)NC2=C(C(=O)NC(S2)C3=CC(=C(C=C3)O)OC)C#N |

**Table S1: Canonical SMILES of top hits**
